# Robust Automated Assessment of Human Blastocyst Quality using Deep Learning

**DOI:** 10.1101/394882

**Authors:** Pegah Khosravi, Ehsan Kazemi, Qiansheng Zhan, Marco Toschi, Jonas E. Malmsten, Cristina Hickman, Marcos Meseguer, Zev Rosenwaks, Olivier Elemento, Nikica Zaninovic, Iman Hajirasouliha

**Affiliations:** Institute for Computational Biomedicine, Department of Physiology and Biophysics, Weill Cornell Medicine of Cornell University, NY, USA; Caryl and Israel Englander Institute for Precision Medicine, The Meyer Cancer Center, Weill Cornell Medicine, NY, USA; Yale Institute for Network Science, Yale University, CT, USA; The Ronald O. Perelman and Claudia Cohen Center for Reproductive Medicine, Weill Cornell Medicine, NY, USA; Institute of Reproduction and Developmental Biology, Imperial College, Hammersmith Campus, London, UK; Instituto Valenciano de Infertilidad, Universidad de Valencia, Valencia, Spain; WorldQuant Initiative for Quantitative Prediction, Weill Cornell Medicine, NY, USA

## Abstract

Morphology assessment has become the standard method for evaluation of embryo quality and selecting human blastocysts for transfer in *in vitro fertilization* (IVF). This process is highly subjective for some embryos and thus prone to human bias. As a result, morphological assessment results may vary extensively between embryologists and in some cases may fail to accurately predict embryo implantation and live birth potential. Here we postulated that an artificial intelligence (AI) approach trained on thousands of embryos can reliably predict embryo quality without human intervention.

To test this hypothesis, we implemented an AI approach based on deep neural networks (DNNs). Our approach called STORK accurately predicts the morphological quality of blastocysts based on raw digital images of embryos with 98% accuracy. These results indicate that a DNN can automatically and accurately grade embryos based on raw images. Using clinical data for 2,182 embryos, we then created a decision tree that integrates clinical parameters such as embryo quality and patient age to identify scenarios associated with increased or decreased pregnancy chance. This IVF data-driven analysis shows that the chance of pregnancy varies from 13.8% to 66.3%.

In conclusion, our AI-driven approach provides a novel way to assess embryo quality and uncovers new, potentially personalized strategies to select embryos with an improved likelihood of pregnancy outcome.

## Introduction

Infertility remains an unremitting reproductive issue that affects about 186 million people worldwide^1^. In the United States, infertility affects approximately 8% of women of child-bearing age^2^. Approximately 44% of women in the U.S. meet the criteria for infertility at a certain point during their reproductive years^3^. Assisted reproductive technology (ART), including in vitro fertilization (IVF), is one of the most common treatments for infertility. IVF involves ovarian stimulation followed by the retrieval of multiple oocytes, fertilization, and embryo culture for 1-6 days in controlled environmental conditions. Embryo quality is then assessed by morphological criteria in an effort to select the best one or two embryos for transfer to the patient’s uterus. Because embryo morphology is only an approximate surrogate for embryo quality, multiple embryos are often transferred, as no method is of high enough accuracy and reliability to assure that a single embryo selected will result in implantation^4^. Indeed, although IVF and embryo-transfer technologies have improved considerably over the past 30 years, the efficacy of IVF continues to remain relatively low^5^.

Conventional embryo evaluation involves observation, assessment, and manual grading of blastocyst morphological features by skilled embryologists. While this selection method is used universally in clinical practice, the evaluation of an embryo based on a static image represents a rather crude, subjective evaluation of embryo quality, which is also time-consuming^6-8^.

Complicating the problem, there continues to be a tendency for inconsistent blastocyst classification, often associated with different grading systems among medical centers. This makes it difficult and challenging to compare selection methodologies and analyze patients undergoing treatments in different clinics. Indeed, attempts to establish a universal grading and selection system have thus far failed to catch on^9^.

Improving the ability to select the embryos with the highest implantation potential would increase pregnancy rates as well as minimize the chance of multiple pregnancies due to the transfer of multiple embryos^10^. Opportunities exist to leverage artificial intelligence (AI) in IVF clinics which have adopted digital imaging as part of their clinical practice, utilizing their time-lapse datasets of many thousands of labeled images. Time-lapse imaging (TLI) is an emerging technology that allows continuous observation of embryo development without removing embryos from controlled and stable incubator conditions^11^. Time-lapse analysis was first used more than three decades ago to study the development of bovine embryos in vitro^12, 13^. Interest in using this technology to assess human embryos has recently grown, as it has been shown to improve selection of the most robust embryos for transfer^14^. This technology also improved IVF cycle outcomes by decreasing the embryos’ exposure to changes in temperature, high oxygen, and fluctuations in pH during culture^15^. In addition, it has enabled embryologists to assess embryo quality by tracking the timing of embryo cleavage events and the temporal intervals between hallmarks observed during embryo development (karyokinesis and cytokinesis)^16^.

Currently, no robust and fully automatic method exists to analyze human embryo data by TLI. Several groups have attempted to use different machine learning approaches for embryo quality analysis, with varying degrees of success^17, 18^ for bovine and mammalian oocytes using artificial neural network (ANN)- and random forest (RF)-based classification, respectively. Their results showed 76.4% (test set = 73 embryos) and 75% (test set = 56 embryos) accuracy for discretization of bovine embryo grades and mammalian oocyte grades, respectively. Furthermore, a few previously published approaches have focused on classifying human embryo quality based on specific features, such as the inner cell mass (ICM) area, trophectoderm (TE) area, zona pellucida (ZP) thickness, and blastocyst area and radius separately^10, 19^. In particular, Filho et al.^19^ presented a semi-automatic grading system of human embryos. They showed that classifiers can have different accuracies for each object (blastocyst extension, ICM, and TE). Their results indicated various accuracy ranges from 67% to 92% for the embryo extension, from 67% to 82% for the ICM, and from 53% to 92% for TE detection; 92% was the highest accuracy achieved across a 73-embryo test set^19^. Although these methods achieved reasonable accuracy in assessing human embryo quality, they require advanced embryological expertise and several preprocessing steps, which are time-consuming.

Deep learning has recently been used to address a number of medical-imaging problems, such as predicting skin lesions or diagnosing disease^20^. Our group has also recently shown that deep learning can significantly improve performance, correctness, and robustness in classifying and quality assessment of digital pathology images in cancer^21^.

In this paper, we introduce a computational method using deep learning techniques (Figure 1) to predict the quality of human embryos. In the first step, our embryologists generated embryo images from TLI and labeled embryos as good-quality or poor-quality. In the second step, a deep neural network (DNN) was trained to automatically assess the quality of the images and evaluated using a blind test set comprising of good- and poor-quality images of human embryos. Finally, a decision tree was used to combine the deep learning-based assessment of embryo quality with clinical data such as patient’s age to identify the (ideal) clinical scenarios associated with a maximized likelihood of pregnancy (Figure 1).

**Figure 1.**
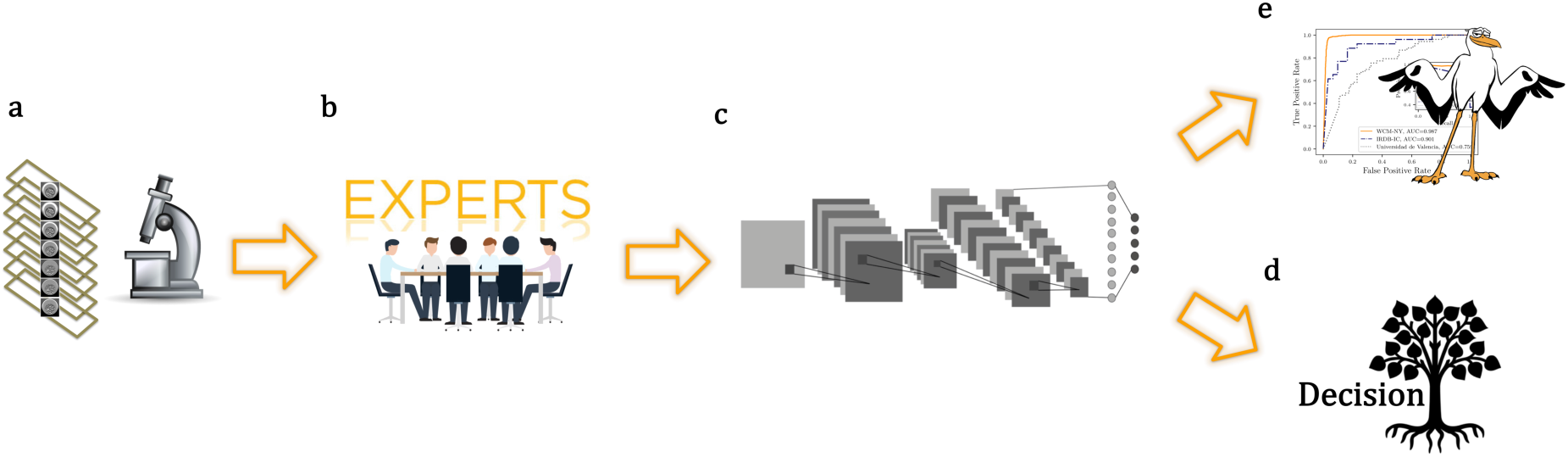
This flowchart demonstrates the design and assessment of STORK. (a) Human embryo images are provided from the embryology lab; (b) the embryo images are labeled by embryologists as good-quality or poor-quality based on their pregnancy likelihood; (c) the labels and clinical information from the extracted images are integrated, and the Inception-V1 algorithm is trained for good-quality and poor-quality classes; (d) the CHAID decision tree is used to investigate the interaction between clinical information, such as patient age with embryo quality; and (e) STORK is evaluated by a blind test set to assess its performance in predicting embryo quality.

## Results

We obtained time-lapse images of 10,148 de-identified embryos, taken at 110 hpi (hours post-insemination) after fertilizing oocytes, at the Center for Reproductive Medicine at Weill Cornell Medicine, New York (WCM-NY). The images were taken at seven focal depths (+45, +30, +15, 0, −15, −30, and −45), constituting a set of 50,392 total images. Trained embryologists evaluated embryo quality using an internal scoring system. To enable the AI analysis, the 10,148 embryos were subsequently classified into three major groups (good-quality = 1,345 embryos, fair-quality = 4,062 embryos, and poor-quality = 4,741 embryos) (Figure 2a) as described in the Methods section.

**Figure 2.**
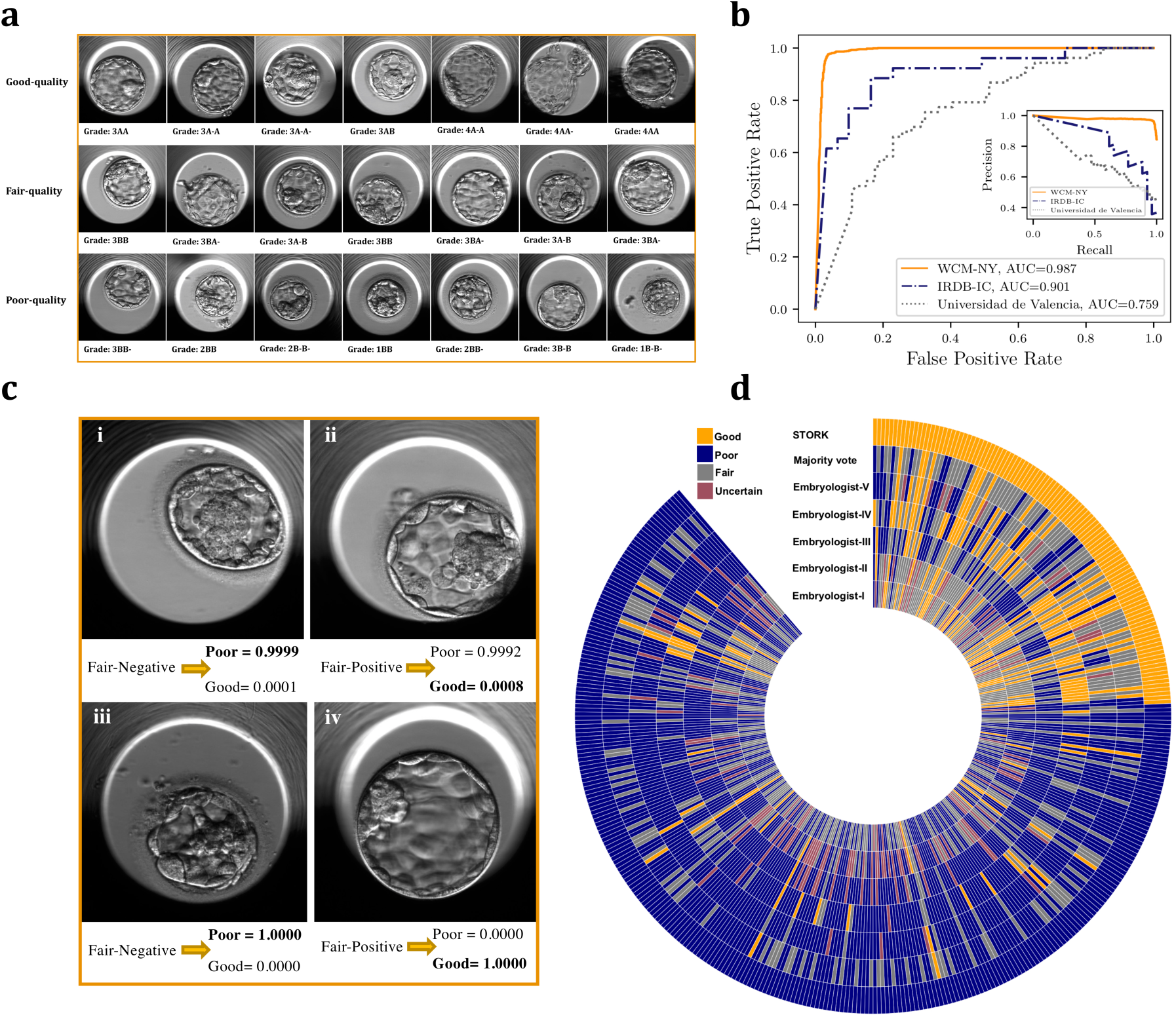
(a) Embryologists evaluate embryo quality using an internal scoring system and subsequently classify them into three major groups (good-quality, fair-quality, poor-quality). (b) Inception-V1 (fine-tuning the parameters for all layers) results for three datasets. WCM-NY: data from the Center for Reproductive Medicine and Infertility at Weill Cornell Medicine of New York; Universidad de Valencia: data from the Institute Valenciano de Infertilidad, Universidad de Valencia; IRDB-IC: data from the Institute of Reproduction and Developmental Biology of Imperial College. (c) STORK classifies the fair-quality images into existing good-quality and poor-quality classes. For example, figures “i” and “ii” are labeled 3A-B according to the Veeck and Zaninovic grading system, while STORK classified them as poor-quality and good-quality, respectively. Also, figures “iii” and “iv” are both labeled 3BB. However, the algorithm correctly classified figure “iii” as poor-quality and figure “iv” as good-quality. As the figure shows, the outcome in the embryos in “ii” and “iv” is positive live birth, whereas it is negative live birth in “i” and “iii”. (d) This circular heatmap demonstrates the agreement between STORK and five embryologists in the labeling of the same images from 394 embryos. The heatmap also compares STORK’s result with the majority vote results from all of the embryologists for 239 embryos. Orange: embryos with good-quality; navy: embryos with poor-quality; gray: embryos with fair-quality; red: embryos that are not labeled due to uncertainty.

We sought to train an Inception-V1 deep learning-based algorithm using the two quality groups at both ends of the spectrum, i.e., good-quality and poor-quality. The Inception-V1 architecture is a transfer learning algorithm, where we initially performed fine-tuning of the parameters for all of the layers. Upon preprocessing and removal of bad-quality images and random selection of a balanced set of images, we were left with a total of 12,001 images with up to seven focal depths (+45, +30, +15, 0, −15, −30, and −45): 6,000 images in 877 good-quality embryos, and 6,001 images in 887 poor-quality embryos. We used 50,000 steps for training the DNN. We then evaluated the performance of STORK using a randomly selected independent test set with 964 good-quality images (141 embryos) and 966 poor-quality embryo images (142 embryos).

### DNN architecture achieves the expert-level classification of embryo images

Our results showed that the trained algorithm was able to identify good-quality and poor-quality images with 96.94% accuracy (1,871 correct predictions out of 1,930 images = 96.94% accuracy) when tested on 964 good-quality and 966 poor-quality embryos.

To measure the accuracy of STORK for individual embryos, we used a simple voting system across multiple image focal depths. If the majority of images from the same embryo were predicted good-quality, then the final quality of the embryo was considered good. For a small number of cases in which the number of good-quality and poor-quality images was equal (e.g., three good-quality and three poor-quality when the number of focal depth was 6), we used STORK’s output probability scores to break the tie. We compared the average STORK probability scores of the good-quality images with the average probability scores of the poor-quality images.

We observed 97.53% accuracy (276 correct predictions out of 283 embryos; Figure 2b) on the blind test set. We also found that training an Inception-V1 model without parameter fine-tuning did not affect performance (accuracy; see Supplementary Figure 1). This observation is in agreement with previous studies using these deep learning techniques^20-22^.

We also found that STORK classified the fair-quality embryo images (4,480 images from 640 embryos) as 82% good-quality (526 embryos) and 18% poor-quality (114 embryos), respectively (Figure 2c). Attesting to the intermediate status of the fair-quality group, the average STORK probability score was 0.98 for good-quality predictions and 0.93 for poor-quality predictions (Supplementary Figure 2).

These STORK probability scores are significantly (p-value <0.01) lower than the probability scores for good-quality and poor-quality images (0.99 on average). Because Inception-V1 was trained for good-quality and poor-quality classes with different pregnancy probabilities (an approximately 58% and 35% chance of pregnancy for good-quality and poor-quality classes, respectively), we wondered if STORK nonetheless produced relevant predictions within the fair-quality class. A closer look showed that embryos with fair-quality images that were classified as poor-quality by STORK had a lower likelihood of positive live birth (50.9%) as compared to those classified as good-quality (61.4% positive live birth, p <0.05 by the two-tailed Fisher’s test).

In addition, we found that fair-quality embryos predicted to be good-quality by STORK came from younger patients (33.98 years old on average) than those predicted to be poor-quality (34.25 years old on average; p-value < 0.01). Interestingly, these numbers are similar to the patients age with good-quality and poor-quality embryos: 33.86 and 34.72 years old on average, respectively. This suggests that STORK finds sufficient structure within embryos classified as fair-quality to make clinically relevant predictions.

### The robustness of STORK

To evaluate STORK’s robustness, we tested its performance by using additional datasets of embryo images obtained from two other IVF centers, Universidad de Valencia and IRDB-IC, comprising 127 (74 good-quality, 53 poor-quality) and 87 (61 good-quality, 26 poor-quality) embryos, respectively (Supplementary Table 3). Our experimental results (See Figure 2b) demonstrate that although the scoring systems used for these centers are different from the system used to train our model, STORK can successfully identify and register score variations and robustly discriminate between them, with an accuracy of 77% (average precision = 0.8, AUC = 0.9) and 70% (average precision = 0.66, AUC = 0.76) for the IRDB-IC and Universidad de Valencia, respectively (Supplementary Table 3).

It is well known that embryo scoring frequently varies among embryologists^23^, mainly due to the subjectivity of the scoring process and different interpretations of embryo quality. We therefore sought to create a small but robust benchmark embryo dataset that would represent the consensus of several embryologists. We asked five embryologists from three different clinics to provide scores for each of 394 embryos generated in different labs. Note that these images were not used in the training phase of our algorithm. The embryo images were scored using the Gardner scoring system^24^ and then mapped onto our simplified three groups (good-quality, fair-quality, and poor-quality; see Supplementary Table 5 and Supplementary Table 2 for the mapping method).

As expected we found a low level of agreement among the embryologists, with only 89 embryos out of the 394 classified as the same quality by all five embryologists (Supplementary Figure 3). Therefore, to create a larger and more accurate gold standard dataset, we used an embryologist majority voting procedure (i.e., the quality of each image was determined by the score given by at least three out of the five embryologists) to classify 239 images (32 good-quality and 207 poor-quality).

When we applied STORK to these 239 images, we found that it predicted the embryologist majority vote with high accuracy (90.4%) and average precision (95.7%). In comparison, STORK agreed with the individual embryologists slightly less often (89.6%, 85.8%, 80.8%, 85.8%, and 88.3% accuracy; 92.1%, 89.5%, 97.4%, 88.3%, and 96.3% average precision). These results indicate that STORK is at least as reliable as any individual embryologist when classifying embryo image quality (Figure 2d).

### Predicting pregnancy likelihood using the trained algorithm for embryo outcome

It is known that factors such as embryo quality, maternal age, the patient’s genetic background, clinical diagnosis, and treatment-related characteristics can affect pregnancy outcome^25, 26^. Because embryo quality is one of the most important of these factors, the ultimate aim of any embryo-assessment approach is to identify embryos that have the highest implantation potential, resulting in live birth^24, 27, 28^.

We explored the possibility of directly predicting the likelihood of pregnancy based on embryo images labeled as “positive” or “negative live birth”. To address this question, we used WCM-NY images associated with 1,620 embryos for which we had the pregnancy outcome (live birth) information (Supplementary Table 1). We allocated 85% of the embryos (1,377 embryos, 9,639 images) to build two classes-“negative live birth” (603 embryos) and “positive live birth” (774 embryos)-as training. There were good- and poor-quality embryos that were assessed by embryologist, in both the “negative live birth” (embryos ‘a’ and ‘b’ in Supplementary Figure 4) and “positive live birth” classes (embryos ‘c’ and ‘d’ in Supplementary Figure 4). Thus, we had embryo images with four different characteristics in two classes (Supplementary Figure 4).

We built a new training algorithm, different from STORK, called DCNN (deep convolutional neural network) to fine-tune the Inception-V1 algorithm using two classes (positive and negative live birth) with 50,000 steps.

Finally, we tested DCNN with 243 randomly selected embryos as a blind test comprising 136 and 107 “positive live birth” and “negative live birth” embryos (1,701 images), respectively (Supplementary Table 1).

We obtained only a 51.85% accuracy for discretization of positive and negative live birth. This suggests that discretization of images based on live birth outcome using embryo morphology alone cannot be useful since other important characteristics, such as the patient’s age and genetic or clinical variations, can affect the pregnancy rate. We refer the reader to Appendix B for a detailed discussion.

Therefore, in the next section we present an alternative method for predicting pregnancy probability based on a state-of-the-art decision tree method that integrates clinical information and embryo quality.

### Decision tree reveals the interaction between clinical information

As we showed in the previous section, embryo quality alone is not enough to accurately determine the pregnancy probability. Therefore, we wondered if we could assess the pregnancy rate by using a combination of embryo quality and patient age, as age is one of the most important clinical variables. For this purpose, we used a hierarchical decision tree method known as a chi-squared automatic interaction detection (CHAID) algorithm^29^.

We designed a CHAID^30, 31^ decision tree using all 2,182 embryos from the WCM-NY database with available clinical information (Figure 3). We then investigated the interaction between patient age (consisting of seven classes: ≤30, 31–32, 33–34, 35–36, 37–38, 39–40, and >41) and embryo quality (consisting of two classes: good-quality and poor-quality), and their effect on live birth outcome. The CHAID algorithm can project interactions between variables and non-linear effects, which are generally missed by traditional statistical techniques. CHAID builds a tree to determine how variables can explain an outcome in a statistically meaningful way^30, 31^. CHAID uses χ^2^ statistics through the identification of optimal multi-way splits, and identifies a set of characteristics (e.g., patient age and embryo quality) that best differentiates individuals based on a categorical outcome (here, live birth) and creates exhaustive and mutually exclusive subgroups of individuals. It chooses the best partition on the basis of statistical significance and uses Bonferroni-adjusted p-values to determine significance with a predetermined minimum size of end nodes. We used a 1% Bonferroni-adjusted p-value, a maximum depth of the tree (n = 5), and a minimum size of end nodes (n = 20) as the stopping criteria. The application of a tree-based algorithm on the embryo data would help to more precisely define the effect of patient age and embryo quality (good-quality or poor-quality) on the live birth outcome, and to better understand any interactions between these two clinical variables (patient age and embryo quality).

**Figure 3.**
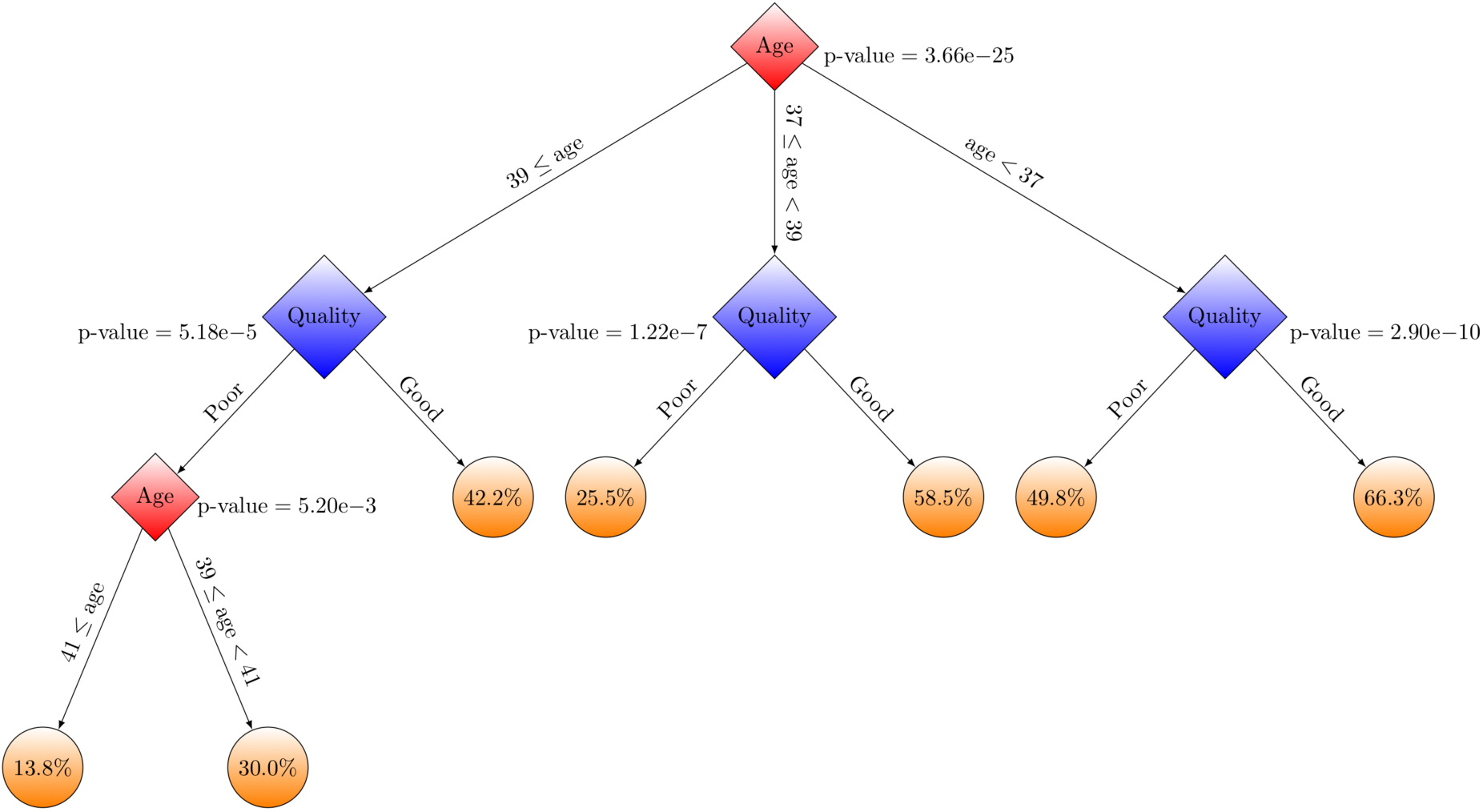
The decision tree shows the interactions between IVF patient ages and embryo quality using CHAID.

Note that while several other classification algorithms could have been employed for the prediction, CHAID was the best fit in terms of model quality criteria, and it enabled a more proper visualization of the decision tree diagram^32, 33^.

As Figure 3 shows, patients were classified into three age groups: (i) ≤36, (ii) 37 and 38, and (iii) ≥39 years old. For each age group, embryos were classified in good- and poor-quality groups.

The results confirm the association between pregnancy probability and patient age. The pregnancy probability for patients with good-quality embryos is significantly (1% Bonferroni-adjusted p-value) higher than that for patients with poor-quality embryos across different ages. Figure 3 indicates that patients ≤36 years old have a higher pregnancy rate compared to patients in the other two age groups. The CHAID decision tree analysis also indicates that the chance of pregnancy varies from 13.8% (e.g., when the embryo is of poor-quality as assessed by STORK and the patient is >41 years old) to 66.3% (e.g., when the embryo is of good-quality and the patient is <37 years old) using IVF (Figure 3).

## Discussion

Computational embryology is a rapidly evolving field. There is enormous potential for using computational approaches to supply prognostic information that cannot be provided by embryologists alone. The STORK framework presented here provides a novel method that can be easily implemented for a wide range of applications, including embryo grading.

Recently, there have been several studies utilizing classical machine learning approaches, such as support vector machine (SVM) and RF, and deep learning methods, such as CNN-basic^17, 18, 34^, for outcome prediction or grade classification. To date, several AI methods have been used to assess blastocysts^35^. Image segmentation and advanced image analysis techniques using neural networks with textured descriptors, level set, phase congruency, and fitting of ellipse methods have been demonstrated in mouse^36^, bovine^17^, and human blastocysts^19, 37^. Studies on human embryos are still very limited and they often involve low numbers of embryos (51–394) from single centers and lack validations in independent cohorts. Furthermore, publications to date have relied on images that were captured using inverted microscopes. However, time-lapse images have the advantage of being consistent in terms of size, lighting, contrast, and quality, and in terms of capturing the timing of embryo development, which is particularly important when quantifying blastocyst expansion.

The aim of this project was to evaluate the utility of DNNs to automatically identify embryo quality. To the best of our knowledge, this is the first study to use higher-level architecture of a DNN algorithm. The advantage of this technique is that instead of only focusing on the predetermined, segmented features that embryologists are trained to analyze, the entire image of the embryo is assessed, allowing for quantification of all the available data. Convolution, therefore, allows the AI to identify patterns in morphological features that we do not know how to assess.

We have demonstrated that deep learning approaches can provide accurate quality assessments in various clinical conditions. Our results show that the accuracy of a DNN primarily depends on the labels that we use to train the algorithm.

Our method yields a cutting-edge sensitivity when performing the challenging task of assessing embryo quality using multi-focal embryo images. Notably, our STORK framework is fully automated and does not require any manual augmentations or pre-processing on the input images. In fact, it provides embryologists or medical technicians a straightforward platform to use without requiring sophisticated computational knowledge. Finally, we designed a decision tree using the CHAID algorithm to investigate the interaction between embryo quality and patient age, and their effect on the pregnancy rate (live birth likelihood).

We also showed that our study raises several important issues regarding embryo clinical conditions, as different deep learning responses could be caused by clinical situations. Further studies are required to clarify the efficiency of the deep learning application in predicting pregnancy outcome.

## Methods

In this section, we present our AI-based method for classifying embryo morphologies. We also discuss how we assessed the accuracy and consistency of the AI classifier in comparison to human classification.

### Embryo images

This study included 10,148 de-identified embryos from our Center for Reproductive Medicine at Weill Cornell Medicine (2012/05 - 2017/12). This study used fully de-identified data and was approved by the Institutional Review Board (IRB) at Weill Cornell Medicine (IRB no. 1401014735). We refer to this dataset as WCM-NY throughout this manuscript. The images were captured using the following technique: EmbryoScope^®^ time-lapse system (Vitrolife, Sweden); built-in microscope: Leica 20x, 0.40 LWD Hoffman modulation contrast objective specialized for 635 nm illumination; camera resolution: 1280*×*1024 pixels, three pixels per µm, monochrome, 8-bit; embryo illumination: 0.032s per image using single red LED (635nm) gives 34µW cm-2 for image acquisition; time between acquisitions: 15-min. cycle time for seven focal planes representing a total of 50,392 images (stored in jpg, 500*×*500 pixels) with about seven focal depths (+45, +30, +15, 0, −15, −30, and −45) captured precisely 110 hpi (Supplementary Figure 5). The standardization of images by the EmbryoScope software was consistent, and the images were labeled using the Veeck and Zaninovic grading system^38^. In addition, these images contain 130 various grades, of which most comprise a few image numbers (Supplementary Table 4). We eliminated from the dataset images that were either very dark or missing an embryo picture, and we selected a balanced set of images for both good-quality and poor-quality classes.

The Veeck and Zaninovic grading system^38^ (Supplementary Table 5) is a slightly modified version of the Gardner system^24^, classifying embryos based on blastocyst expansion (grades 1 to 6), cell abundance, and conformity in the ICM (grades A, B, and C) and TE (grades A, B, and C) (Supplementary Table 5). In addition to our WCM-NY data, we used two other datasets from the Universidad de Valencia and the Institute of Reproduction and Developmental Biology of Imperial College (IRDB-IC). The data from the Universidad de Valencia was graded based on a slightly different of the Gardner scoring system known as Asebir^39^. Compared to the Gardner system, Asebir uses five rather than six expansion categories and changes the ICM and TE rating terminology to single A, B, C, and D letters (Supplementary Table 5). The IRDB-IC data was graded using the Gardner scoring system.

### Classification and diagnostic framework

This study presents a framework (see Figure 1) to classify different embryo images based on Veeck and Zaninovic grades (Supplementary Table 4) and map those grades to good- and poor-quality blastocyst grades. Here, we used the WCM-NY embryos and clinical information from a subset of these embryos, such as grades and patient age.

We divided the images into training, validation, and test groups. We allocated 70% of the images to the training group and the remaining 30% to the validation and test groups (Supplementary Table 1). The training, validation, and test sets did not overlap.

### Algorithm architectures and training methods

We employed a DNN for embryo image analysis based on Google’s Inception-V1^40^ architecture, which offers a very effective run-time and computational cost^41, 42^. To train this architecture, we used transfer learning. We employed a pre-trained network and fine-tuned all outer layers^43^ using the WCM-NY images. We also compared this transfer learning approach to training the network from scratch.

### Evaluation of method and implementation details

To implement the STORK framework, we used the Tensorflow version 1.4.0^44^ and the Python library TF-Slim for defining, training, and evaluating models in TensorFlow. All training of our deep learning methods were performed on a server running the SMP Linux operating system. This server is powered by four NVIDIA GeForce GTX 1080 GPUS with 8 GB of memory for each GPU and 12 1.7-GHz Intel Xeon CPUs.

To evaluate the performance of our methods, we used an *accuracy* measure, which is the fraction of correctly identified images^21^. The accuracy is formally defined as Tnu /(TNu + FNu), where TNu (true number) and FNu (false number) are the number of correctly and incorrectly classified images.

To assess the performance of different algorithms, precision-recall curves (PRCs) were used. Here, precisions and recalls are presented by average for multi-class datasets. Additionally, receiver operating characteristics (ROCs) were estimated. The ROC curve is depicted by plotting the true positive rate (TPR) versus the false positive rate (FPR) at various threshold settings. The accuracy is measured by the area under the ROC curve (AUC)^45, 46^.

## Acknowledgements

We acknowledge Dr. Fabien Campagne for useful discussions and providing additional computing resources for our analysis. This work was supported by start-up funds (Weill Cornell Medicine) to IH. EK was supported by Swiss National Science Foundation under grant number 168574.

## Author contributions statement

PK, EK, JEM, CH, MM, NZ, OE, and IH conceived the study. PK, EK, OE, and IH conceived the method and designed the algorithmic techniques. QZ, MT, CH, MM, and NZ generated the datasets and prepared and labeled the images for various grades. PK and EK wrote the codes and performed computational analysis with input from OE and IH. ZR provided critical reading and suggestions. PK, EK, QZ, OE, NZ, and IH wrote the paper, and all authors read, edited, and approved the final manuscript.

## Supplementary Information

## Appendix A: Embryologists split and merge the quantity grades

In this project, skilled embryologists determined the quantitative scores based on the grading system of Veeck and Zaninovic^38^. This grading system has three components: The first is a number showing the level of blastocyst expansion (CM, 1, 2, 3, 4, and 5), the second is a letter indicating the cell abundance and conformity in the ICM (grades A, B, C, and D), and the third is a letter quantifying the quality of TE cells (grades A, B, C, and D), which are extra-embryonic tissues that support the embryo proper (see Supplementary Table 5).

For the first step of this project, the embryologists selected 13,931 images of embryos with good- and poor-quality based on their pregnancy outcome. The embryologists labeled the embryo images to map certain quantitative scores from the grading system of Veeck and Zaninovic (e.g., 1BB vs. 3AA) to just two quality grades: good-quality and poor-quality (Supplementary Table 5). In this regard, any score that contained B- or C and an extension rate equal to or less than three was considered part of the poor-quality group (<35% pregnancy chance). In addition, any score with two A or A-grades, or one A with B, with an extension of 3 or greater could be labeled as good-quality (>58% pregnancy chance). However, the experts debated about some scores (e.g., 3BB, 3BA-), putting them in a separate category (fair-quality) or classifying them as good-quality, as their pregnancy likelihood was about 48–50%. The complete list of scores and their quality map are shown in Supplementary Table 2. In total, 86 out of 130 scores had images with clinical information, and 84 scores contained a small number of images in their cohorts (Supplementary Table 4).

We converted various quantitative grades related to other data resources to the Veeck and Zaninovic^38^ scoring system before testing our trained algorithm with other clinical resources (Supplementary Table 3). For instance, the 3AA grade in our WCM-NY dataset is equivalent to the BEaa grade in the Universidad de Valencia dataset and the 4AA grade in the IRDB-IC dataset (Supplementary Table 5), which is based on the Gardner system^9, 24^. Notably, these two datasets are less accurate compared to the WCM-NY dataset due to variations in the grading systems. Information about the grading systems used for the different datasets is shown in Supplementary Table 5.

## Appendix B: Predicting pregnancy rate based on morphological quality of embryos

We wondered what explained the low accuracy of DCNN in predicting pregnancy rate via positive and negative live birth. To find the reason, we looked closer at the results for embryos with four different characteristics (Supplementary Figure 4) that we integrated into two classes (positive and negative live birth).

We found that 28.85%, 47.27%, 41.02%, and 71.13% accuracy for a randomly selected test set (243 embryos) comprised “negative live birth” with “good-quality” (52 embryos) (embryo ‘a’ in Supplementary Figure 4), “negative live birth” with “poor-quality” (55 embryos) (embryo ‘b’ in Supplementary Figure 4), “positive live birth” with “poor-quality” (39 embryos) (embryo ‘c’ in Supplementary Figure 4), and “positive live birth” with “good quality” (97 embryos) (embryo ‘d’ in Supplementary Figure 4), respectively. This suggests that the trained algorithm can classify images based only on their quality (good or poor) while disregarding their outcome (positive or negative live birth) (Supplementary Figure 4). Therefore, the accuracy of DCNN could be increased if we utilized a larger number of images with “poor-quality and negative live birth” and “good-quality and positive live birth” in our test set. Moreover, the DCNN performance decreased due to the integration of good- and poor-quality images with, for example, “negative live birth” in a single class (e.g., embryos ‘a’ and ‘b’ in Supplementary Figure 4).

---

**Supplementary Table 1.** Four datasets showing different images (different number of embryos and clinical information) selected from the databases of WCM-NY (three datasets) and the Universidad de Valencia (one dataset) to assess the performance of STORK across different conditions.

| Datasets | Dataset representation | Labels of inputs and outputs | Number of classes and images |
| --- | --- | --- | --- |
| Good-Poor | 110hpi images of embryos (WCM-NY) | Discrimination of good- and poor-quality of embryos | 2 classes: 12,001 images for training and 1,930 images for test set |
| Outcome-Quality | 110hpi images of embryos (WCM-NY) | Discrimination of positive and negative outcome of embryos through good- and poor-quality of embryos | 2 classes: 9,639 images for training and 1,701 images for test set |
| Five-Experts | 110hpi images of embryos (WCM-NY and Universidad de Valencia) | Discrimination of good- and poor-quality of embryos | 2 classes: 12,001 images for training (STORK as trained algorithm by WCM-NY dataset) and 394 embryos for test set from Universidad de Valencia database |

**Supplementary Table 2.** The quantity scores that the algorithm is trained for. The embryologists categorized the scores into two groups (classes) and labeled them as good-quality and poor-quality.

| The list of grades | The quality map |
| --- | --- |
| 3-4AA, 4A-A, 4A-A-, 5AA-, 4AB, 5A-A-, 4AA-, 4AA, 3A-A, 3AA, 3AA-, 3AB, 3A-A- | good-quality |
| 1-2B-/CB, 1-2B-/CB-/C, 1-2B-C, 1B-/CB-/C, 1BC, 1CB-/C, 1CC, 2-2B-C, 2-3BC, 3B-B-/B, 3BC, 3CA-, 3CB, 3CB-, 3CC, 1-2B-/CB-, 1B-/CB, 1B-/CB-, 1B-/CC, 2-3B-/CB, 2-3B-/CB-, 2B-/CB, 3B-/CB-/C, 3B-/CC, 1-2BB-/C, 2B-/CB-/C, 3B-C, 1BB-/C, 1BB-/C, 1-2B-B-/C, 1B-C, 2-3BB-/C, 2-3B-B-/C, 2B-/CB-, 3B-/CB-, 3B-/CB, 2BB-/C, 1B-B-/C, 2B-B-/C, 3BB-/C, 3B-B-/C, 1-2B-B, 1-2B-B-, 2-3B-B-, 1-2BB-, 2-3B-B, 1B-B, 1BB-, 2-3BB-, 1-2BB, 1B-B-, 2B-B, 2B-B-, 2BB-, 3B-B-, 1BB, 2BB, 3B-B, 3BB- | poor-quality |

**Supplementary Table 3.** The results of applying STORK on various datasets to discriminate two classes of embryo quality. WCM-NY: The Center for Reproductive Medicine and Infertility at Weill Cornell Medicine of New York; Universidad de Valencia: Institute Valenciano de Infertilidad, Universidad de Valencia; IRDB-IC: Institute of Reproduction and Developmental Biology of Imperial College.

| Datasets | Grades | Number of test embryos | STORK accuracy |
| --- | --- | --- | --- |
| WCM-NY | 3AA, 3AA-, 3AB, 3A-A-, 5AA-, 3A-A, 3-4AA, 4AA (good-quality), 3BB-, 2BB-, 2BB, 1BB, 1B-B, 2B-B-, 1BB-, 1B-B-, 2B-B, 1-2BB, 3B-B, 1-2B-B-, 1-2BB-, 1BB-C, 1-2B-B, 3CB (poor-quality) | 283 | 97.53% |
| Universidad de Valencia | BEab, BEaa, BHiaa, BHiab, BHab (good-quality), BCbb, BCbc, BEcc, BEbc, BCcc, BEcb, BCcb (poor-quality) | 127 | 70.08% |
| IRDB-IC | 4Aa, 4Ab, 5Ab, 5Aa (good-quality), 2Cb, 4Bc, 2Bc, 4Cc, 3Bc, 2Cc, 1Bb, 3Cc (poor-quality) | 87 | 77.01% |

**Supplementary Table 4.** Characteristics of 130 various grades and their image numbers.

| The morphological grades | Class size |
| --- | --- |
| 1-2B-/CB, 1-2B-/CB-/C, 1-2B-C, 1-2BA-, 1B-/CB-/C, 1BC, 1CB-/C, 1CC, 2-2B-C, 2-3B-A-, 2-3BA, 2-3BC, 2AA, 2AB-/C, 3-4AA, 3A-B-/C, 3B-/CA, 3B-B-/B, 3BC, 3CA-, 3CB, 3CB-, 3CC, 4A-A, 4A-A-, 4AB-, 4B-/CB, 4B-/CB-, 5A-B, 5AA-, 5BA, 5BA-, 5BB-, 1-2A-B-, 1-2B-/CB-, 1B-/CB, 1B-/CB-, 1B-/CC, 2-3AA, 2-3AA-, 2-3B-/CB, 2-3B-/CB-, 2A-B-/C, 2B-/CB, 2BA, 3A-C, 3B-/CB-/C, 3B-/CC, 4B-B-, 4BB-, 5B-B, 1-2BB-/C, 1AB, 2B-/CB-/C, 3B-C, 4AB, 4B-B, 5A-A-, 5BB, 6BB, 1BB-/C, 1BB-/C, 1-2B-B-/C, 1A-B-, 1B-C, 2-3AB-, 2-3BB-/C, 4A-B, 4AA-, 4BA-, 2-3B-B-/C, 2A-A-, 2B-/CB-, 3B-A, 4AA, 2-3BA-, 1-2A-B, 1A-B, 2BA-, 3B-/CB-, 2AB, 5B-B-, 2AB-, MOR | Less than 10 images per grade |
| 2-3A-B-, 3B-/CB, 2A-B-, 2BB-/C, 3B-A-, 4BB, 2-3AB, 2-3A-A-, CM, CAVM, 1B-B-/C, 2B-B-/C, 3BB-/C, 3BA, 3B-B-/C, 2A-B, 1-2B-B | More than 10 and less than 50 images per grade |
| 2-3A-B, 1-2B-B-, 2-3B-B-, 3A-A, 1-2BB-, 3AB-, 2-3B-B | More than 50 and less than 100 images per grade |
| 1B-B, 1BB-, 2-3BB-, 1-2BB, 1B-B-, 3A-B-, 3AA, 2B-B, 3BA-, 3AA-, 2B-B-, 2-3BB, 3AB, 2BB-, 3B-B-, 1BB, 3A-A- | More than 100 and less than 500 images per grade |
| 2BB, 3B-B, 3BB-, 3A-B, 3BB | More than 500 images per grade |

**Supplementary Table 5.** Information about different grading systems in three clinics.

| Objects | Veeck and Zaninovic | Gardner | Asebir |
| --- | --- | --- | --- |
| Expansion |  |  |  |
|  | CM | 1 | BT |
|  | 1 | 2 | BT |
|  | 2 | 3 | BC |
|  | 3 | 4 | BE |
|  | 4 | 5 | BHi |
|  | 5 | 6 | BH |
| ICM |  |  |  |
|  | A | A | A |
|  | B | A/B | B |
|  | C | B/C | C |
|  | D | C | D |
| TE |  |  |  |
|  | A | A | A |
|  | B | A/B | B |
|  | C | B/C | C |
|  | D | C | D |

**Supplementary Figure 1.**
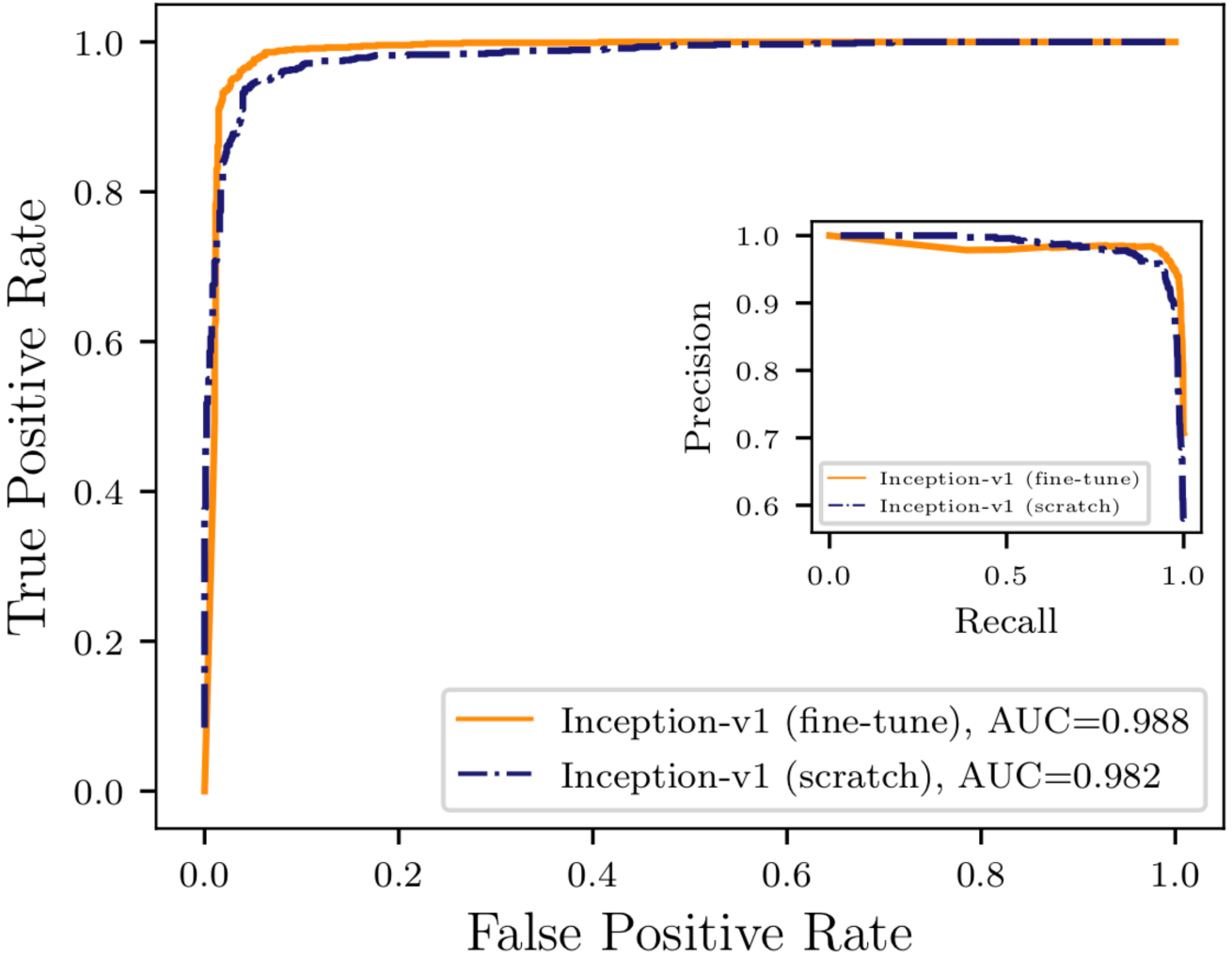
Inception-V1 via two different training methods (fine-tuning the parameters for all layers and training from scratch) in good-quality and poor-quality embryo quality discrimination dataset.

**Supplementary Figure 2.**
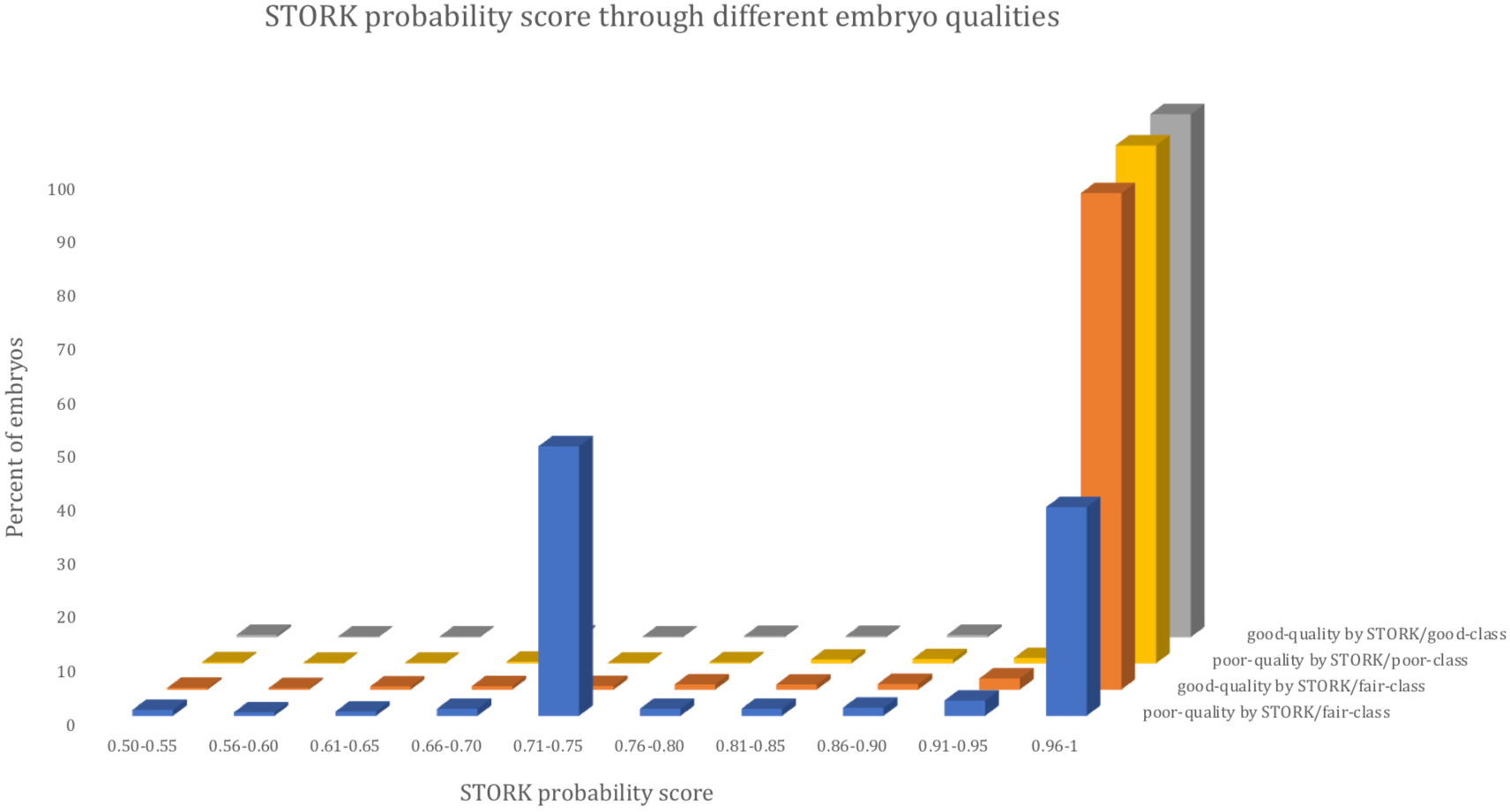
STORK gives each embryo in the fair-quality class a probability score and classifies them into two groups: good- and poor-quality. While the score for embryos that are relabeled by STORK as good-quality and poor-qulity is 0.98 for good-quality and 0.93 for poor-quality, the average probability score for both good-quality and poor-quality classes labeled by embryologists as good-quality and poor-quality is 0.99.

**Supplementary Figure 3.**
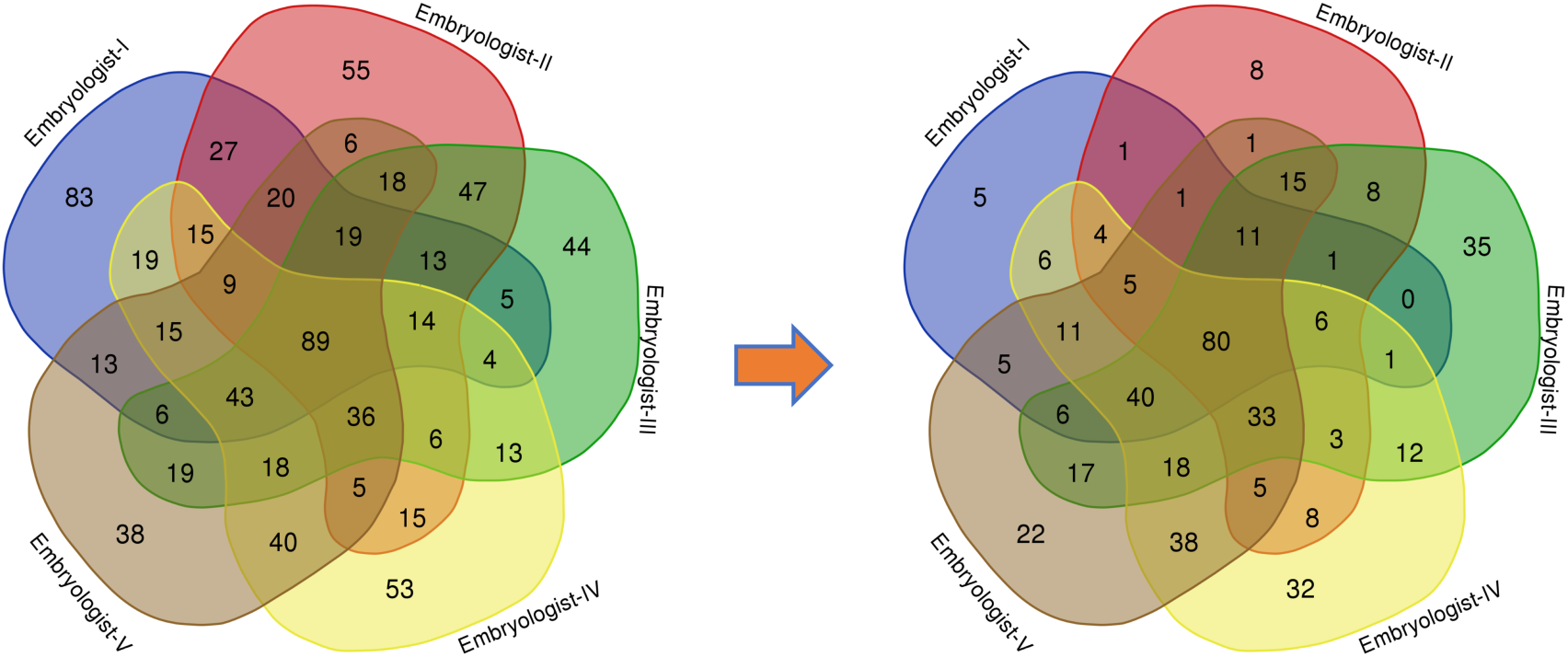
This diagram^47^ demonstrates the agreement among embryologists (Venn diagram on left) and the agreement between STORK and five embryologists (Venn diagram on right) in the labeling of the same embryo images. The colors indicate different embryologists, and the numbers represent the number of embryos.

**Supplementary Figure 4.**
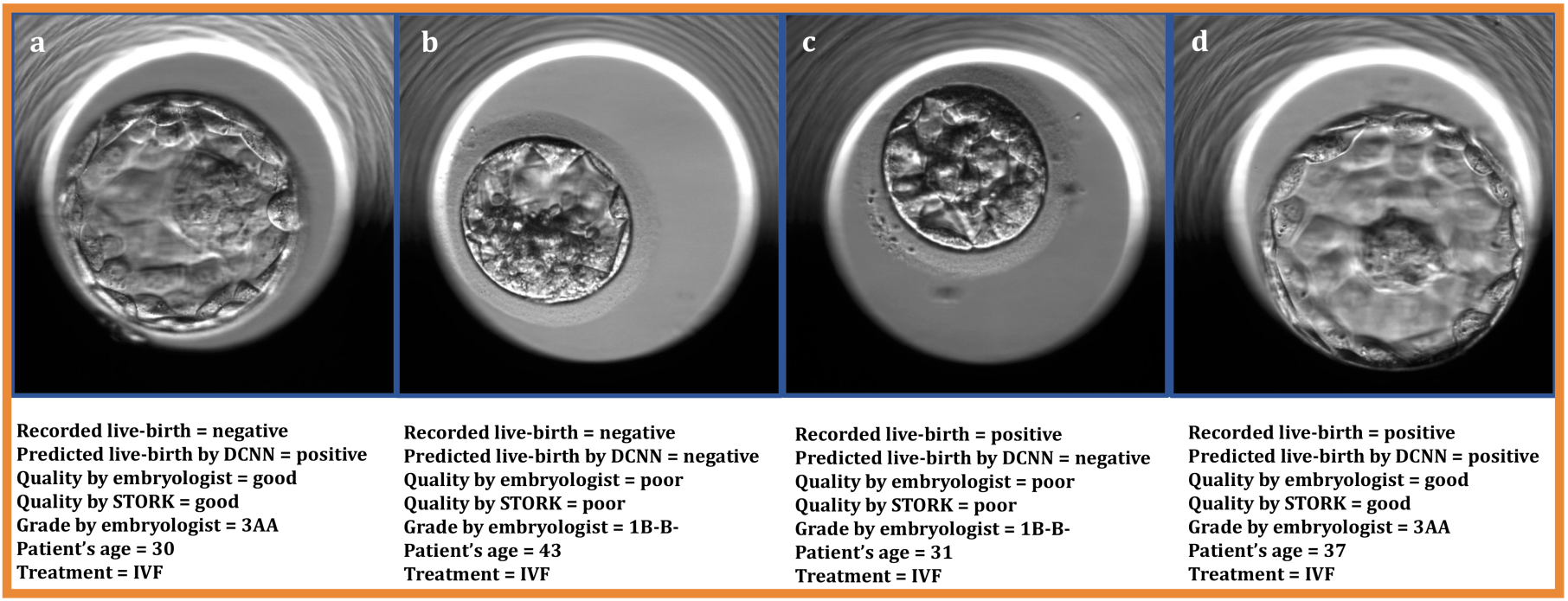
The DCNN classifies embryo images with positive and negative live birth labels with a focus on their morphological quality. For example, embryos “a” and “d” are recorded by the laboratory data manager as negative live birth and positive live birth, respectively. DCNN, however, predicted positive live birth for embryos “a” and “d” because they both have good morphological quality. Embryos “b” and “c” are recorded as negative live birth and positive live birth, respectively. However, the algorithm again classified both embryos “b” and “c” as negative live birth because they have poor-quality.

**Supplementary Figure 5.**
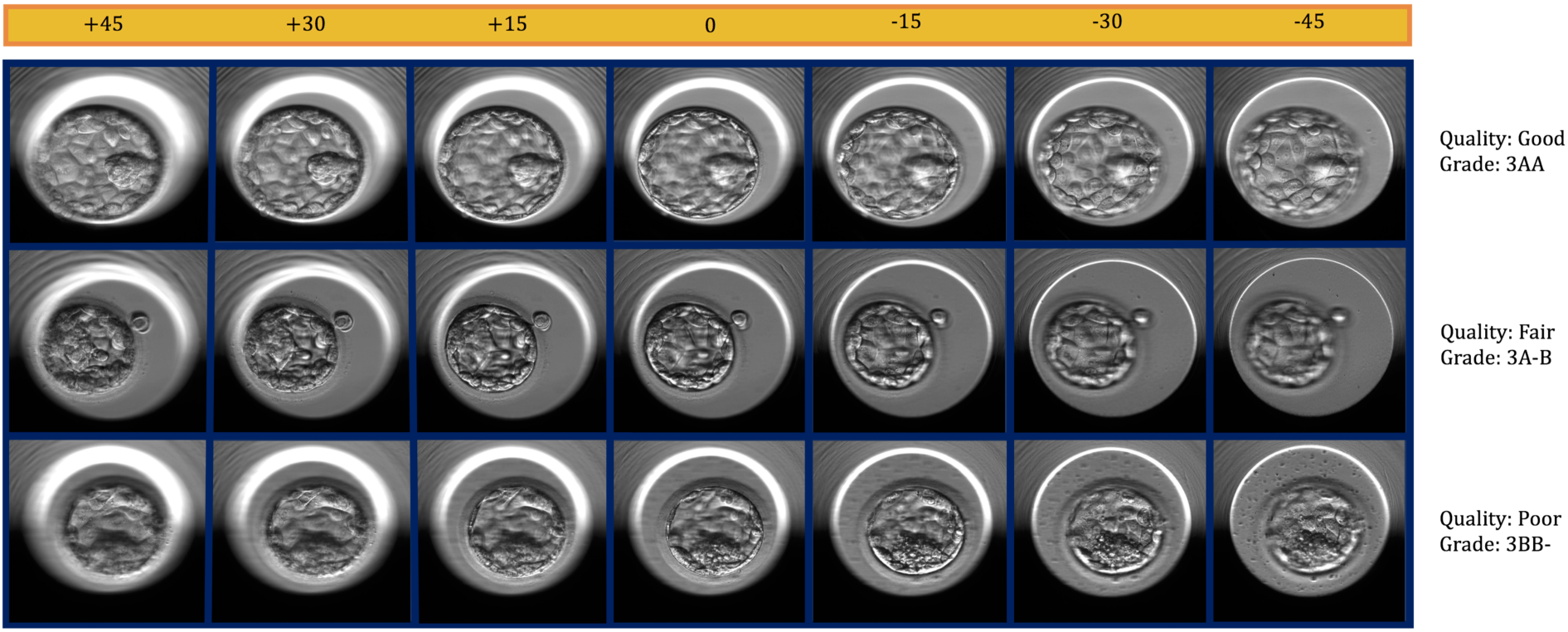
This figure shows three examples of Veeck and Zaninovich grades and their corresponding quality labels across seven focal depths.

